# Inhibition of CAMKK2 impairs autophagy and castration-resistant prostate cancer via suppression of AMPK-ULK1 signaling

**DOI:** 10.1101/2020.06.02.130088

**Authors:** Chenchu Lin, Alicia M. Blessing, Thomas L. Pulliam, Yan Shi, Sandi R. Wilkenfeld, Jenny J. Han, Mollianne M. Murray, Alexander H. Pham, Kevin Duong, Sonja N. Brun, Reuben J. Shaw, Michael M. Ittmann, Daniel E. Frigo

**Author notes:** Corresponding author: Daniel E. Frigo, Department of Cancer Systems Imaging, The University of Texas MD Anderson Cancer Center, 1881 East Road, 3SCR4.3618 – Unit 1907, Houston, TX 77054. These authors contributed equally to this work (Chenchu Lin, Alicia M. Blessing).

## Abstract

Previous work has suggested androgen receptor (AR) signaling mediates cancer progression in part through the modulation of autophagy. Accordingly, we demonstrate that chloroquine, an inhibitor of autophagy, can inhibit tumor growth in preclinical mouse models of castration-resistant prostate cancer (CRPC). However, clinical trials testing chloroquine derivatives in men with CRPC have yet to yield promising results, potentially due to side effects. We hypothesized that identification of the upstream activators of autophagy in prostate cancer could highlight alternative, context-dependent targets for blocking this important cellular process during disease progression. Here, we used molecular (inducible overexpression and shRNA-mediated knockdown), genetic (CRISPR/Cas9), and pharmacological approaches to elucidate an AR-mediated autophagy cascade involving Ca^2+^/calmodulin-dependent protein kinase kinase 2 (CAMKK2; a kinase with a restricted expression profile), 5’-AMP-activated protein kinase (AMPK) and Unc-51 like autophagy activating kinase 1 (ULK1). These findings are consistent with data indicating CAMKK2-AMPK-ULK1 signaling correlates with disease progression in genetic mouse models and patient tumor samples. Importantly, *CAMKK2* disruption impaired tumor growth and prolonged survival in multiple CRPC preclinical mouse models. Finally, we demonstrate that, similar to CAMKK2 inhibition, a recently described inhibitor of AMPK-ULK1 signaling blocked autophagy, cell growth and colony formation in prostate cancer cells. Taken together, our findings converge to demonstrate that AR signaling can co-opt the CAMKK2-AMPK-ULK1 signaling cascade to promote prostate cancer by increasing autophagy. Further, we propose that an inhibitor of this signaling cascade could serve as an alternative, more specific therapeutic compared to existing inhibitors of autophagy that, to date, have demonstrated limited efficacy in clinical trials due to their toxicity and poor pharmacokinetics.

## Introduction

Prostate cancer is the second leading cause of cancer mortality among men in the United States(1). While most prostate cancers can be treated effectively with surgery and/or radiation, a significant number of men present with *de novo* metastatic disease or progress following initial treatment. The standard of care for advanced prostate cancer is androgen deprivation therapy (ADT) due to the central role of the androgen receptor (AR) in almost all prostate cancers(2). Although ADT is initially effective in slowing the cancer, it invariably fails within 2-3 years, after which the disease progresses to a stage referred to as castration-resistant prostate cancer (CRPC). There is currently no cure for CRPC. Interestingly, despite the failure of ADT in CRPC, the overwhelming majority of prostate cancers are still driven by AR as a result of a variety of AR reactivation mechanisms (ex. increased intratumoral androgen synthesis, *AR* gene and enhancer amplifications, splice variants, etc)(2). As such, AR and processes downstream of the receptor remain viable therapeutic targets in CRPC.

In an effort to identify downstream effectors of AR signaling in prostate cancer, we demonstrated *CAMKK2*, encoding the Ca^2+^/calmodulin-dependent protein kinase kinase 2 (CAMKK2) protein, to be a direct AR target gene in prostate cancer(3). These data were soon validated by several other groups(4-6). CAMKK2 expression correlated with both initial response to ADT and transition to CRPC in multiple clinical cohort tissue microarrays (TMAs)(4). In addition, CAMKK2 tracked with Gleason grade and was elevated in different genetically engineered mouse models (GEMMs) of prostate cancer(5, 7). The specific AR binding site we first identified that regulates *CAMKK2* expression(3) was later confirmed by others and shown to be one of the most robust AR binding sites in CRPC patient samples(6). Functionally, CAMKK2 is required for maximum AR-mediated prostate cancer cell growth, migration and invasion in cell culture and tumor growth in xenograft and GEMMs(3-5, 7, 8).

Androgens, in a CAMKK2-dependent manner, increased the phosphorylation of AMP-activated protein kinase (AMPK) on threonine-172 of its α catalytic subunit’s activation loop. Threonine-172 p-AMPK levels correlated with prostate cancer relative to benign prostatic tissue and were further elevated in biochemically recurrent disease(9). Importantly, we previously demonstrated that many of the pro-cancer effects of CAMKK2 in prostate cancer are mediated through the activation of AMPK(3). Accordingly, knockdown of AMPK impaired AR-mediated prostate cancer cell growth(9). These data indicate that AR-CAMKK2 signaling can promote prostate cancer in part through AMPK, a known regulator of macroautophagy(10).

Macroautophagy, herein referred to as autophagy, is a highly conserved process whereby cellular components are captured and delivered to a double membrane vesicle known as an autophagosome, and subsequently degraded by the lysosomal system(11). Autophagy can function as a survival mechanism in response to stress by recycling the lysosomal breakdown products towards essential processes. Autophagy can also serve as a cellular quality control mechanism by removing damaged organelles and toxins. Therefore, autophagy is of importance in physiological processes as well as diseases such as cancer(12). However, the role of autophagy in cancer is complicated and context dependent(13-18). For example, autophagy can protect cells and tissues from damage and impair malignant transformation(19, 20). Conversely, in more advanced cancers, autophagy can enable cells to evade apoptosis in hypoxic and nutrient-deficient environments as well as promote drug resistance(21-24). In prostate cancer, studies from our laboratory and others using cell lines, xenografts and genetic mouse models indicate that autophagy can promote disease progression(25-32). These preclinical data provided the rationale for a series of clinical trials (NCT04011410, NCT00726596, NCT00786682, NCT03513211, NCT01828476, NCT02421575, NCT01480154) that tested the efficacy of chloroquine derivatives such as hydroxychloroquine in men with prostate cancer(33). Chloroquine and hydroxychloroquine were chosen because they 1) are already FDA-approved for the treatment of malaria and rheumatological disorders and 2) have been demonstrated to impair autophagic flux by increasing lysosomal pH and decreasing autophagosome-lysosome fusion(34, 35). Hence, chloroquine and hydroxychloroquine represented potential clinical grade inhibitors of autophagy that could be rapidly repurposed for the treatment of cancer. To date, however, these trials, as well as similar trials in other tumors types, have yielded mixed results(15). To that end, a major challenge has been achieving high enough concentrations of chloroquine or hydroxychloroquine in patients to consistently block autophagy without major side effects(16). The chloroquine-mediated side effects may in part be due to the mechanism of action of this drug. Chloroquine-like compounds inhibit autophagy by blocking the late lysosomal step. Since lysosomal function is required for processes beyond autophagy, this indicates that chloroquine is not specific for autophagy. Hence, we speculate that targeting other steps in autophagy could provide an improved therapeutic window.

We previously demonstrated that androgens, in an AR-dependent manner could increase autophagy and autophagic flux through multiple mechanisms(25, 26). These included indirect activation of autophagy through increases in reactive oxygen species (ROS) and the expression of several core components of the autophagic machinery. Given AMPK’s known link to autophagy(36-38) and our previous findings that AR could increase AMPK activity in a CAMKK2-dependent manner(3, 8), we sought to determine whether the increased CAMKK2 observed in AR+ prostate cancer could be driving disease progression in part through activating autophagy. We also reasoned that delineation of this signaling cascade, combined with the restricted expression profile of CAMKK2 and tolerance for its systemic inhibition in mice(39, 40), could nominate alternative ways to safely block autophagy in men with prostate cancer.

## Results

### Chloroquine impairs CRPC xenograft growth

To initially assess the effect of chloroquine in a preclinical model of CRPC, castrated NSG mice were injected with CRPC 22Rv1 cells stably expressing firefly luciferase (22Rv1-fLuc). When tumors became palpable, mice were randomized to PBS/control and chloroquine treatment groups (Fig. 1A). The tumors were monitored by bioluminescence imaging (BLI) and caliper measurements until the maximum allowable size. In the first two weeks, the BLI clearly showed inhibition of tumor growth by chloroquine (Fig. 1B). However, the bioluminescence intensity lost its sensitivity once the tumors grew large (data not shown), which likely resulted from a lack of oxygen or necrosis in the center of the large tumors. Despite this, the tumor volume demonstrated that chloroquine treatment decreased tumor growth rate (Fig. 1C). The decreased tumor growth rate corresponded to a prolonged survival (Fig. 1D). Compared to the vehicle group, fewer cell nuclei were stained by hematoxylin (Fig. 1E). The reduced tumor growth appeared to be a product of reduced proliferation and increased apoptosis as assessed by BrdU and TUNEL staining, respectively (Fig. 1E). These observations suggest that inhibiting autophagy using chloroquine can reduce proliferation and increase apoptosis, ultimately decreasing CRPC growth and prolonging survival.

**Fig 1.**
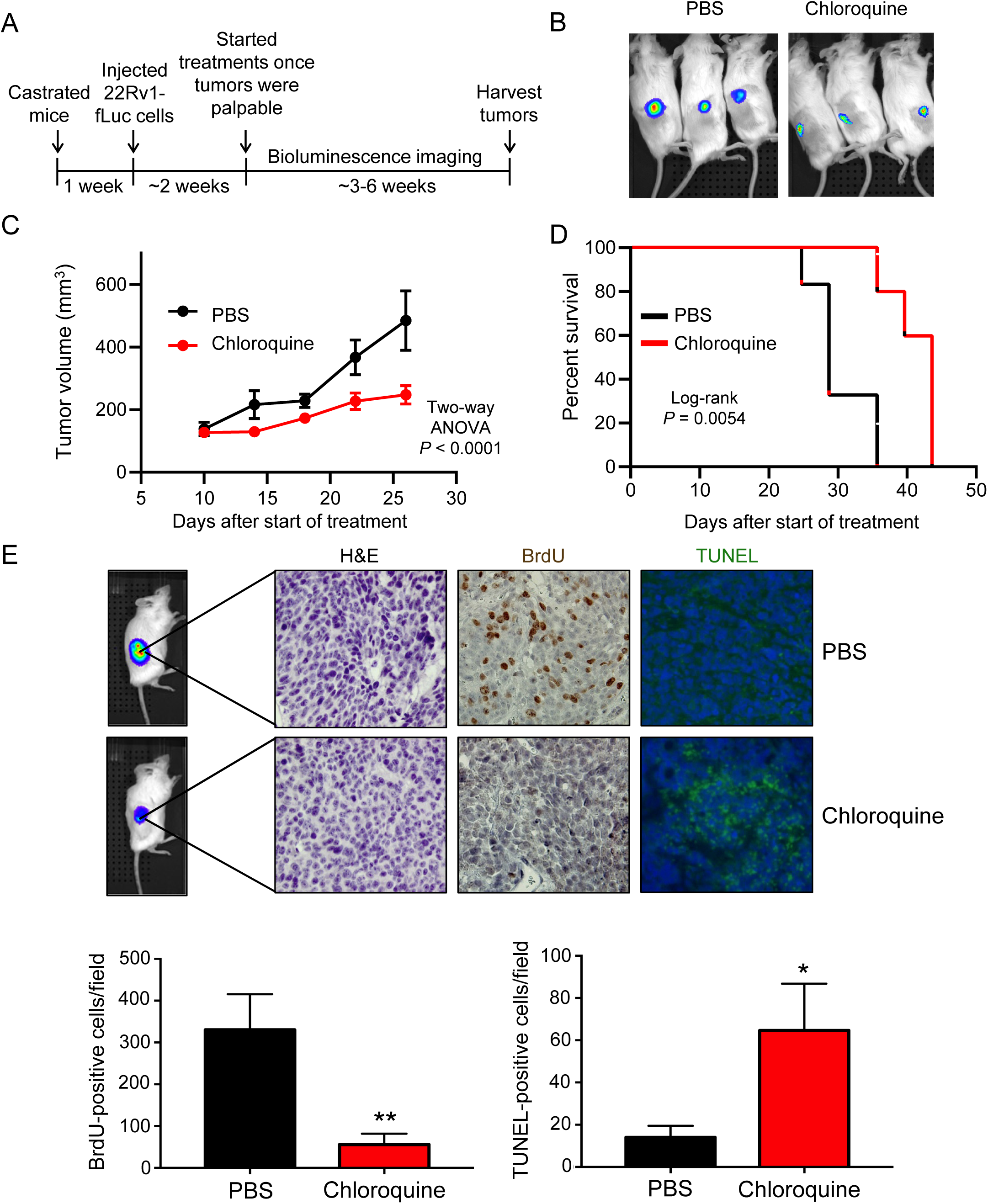
Chloroquine inhibits castration-resistant prostate cancer (CRPC) growth *in vivo.* (A) Schematic of xenograft study using CRPC 22Rv1-fLuc cells in castrated NSG mice treated via intraperitoneal injections (IP) once/day, 6 days/week with vehicle (PBS) or 60 mg/kg/day chloroquine (PBS: n=6, chloroquine: n=7). (B) Bioluminescence imaging of six representative mice bearing tumors. PBS = vehicle. (C) Tumor growth curves of 22Rv1-fLuc xenograft mice treated with vehicle (PBS) or chloroquine. *P* values were calculated using two-way ANOVA. (D) Kaplan-Meier survival curve of 22Rv1-luc xenograft mice following chloroquine treatment. *P* value was calculated using log-rank test. (E) H&E, BrdU and TUNEL staining in the xenograft tumors (*top*). Quantification of BrdU and TUNEL staining (*bottom*). *P* values were calculated using two-tailed *t* test. \**P* < 0.05, \*\**P* < 0.01.

### CAMKK2 promotes autophagy and autophagic flux in prostate cancer

Although chloroquine derivatives have been tested in cancer clinical trials, the high dosage needed in patients to maintain autophagy inhibition remains a challenge that limits the therapeutic window of this class of compounds. We propose that targeting upstream regulators of autophagy may offer safer, alternative options for inhibiting autophagy. *CAMKK2* has previously been shown to be a direct transcriptional target of AR in prostate cancer that promotes the phosphorylation and activation of AMPK(3, 9). Given the critical role of AMPK in autophagy(10, 37, 38, 41), we investigated whether CAMKK2 augmented autophagy in prostate cancer. To do this, we first engineered hormone-sensitive LNCaP cells to inducibly express CAMKK2 in the presence of doxycycline (DOX) (LNCaP-*CAMKK2*). We then examined via immunoblot the effect of CAMKK2 overexpression on AMPK phosphorylation and the accumulation of phosphatidylethanolamine-conjugated LC3B (LC3BII), a marker of autophagy (Fig. 2A). CAMKK2 overexpression increased p-AMPK and conversion of LC3BI to LC3BII (Fig. 2A). Likewise, CAMKK2 overexpression increased GFP-LC3 puncta formation, indicative of increased autophagosome formation (Fig. 2B). To further confirm the effects on autophagy, transmission electron microscopy (TEM) was used to verify the increased number of autophagic vesicles (autophagosomes and autophagolysosomes) following CAMKK2 expression (Fig. 2C). Given the high expression of CAMKK2 in AR+ CRPC(3, 4, 6, 42), we next knocked out *CAMKK2* in C4-2 cells, an LNCaP-derived CRPC model, using CRISPR-Cas9 to assess the effects of *CAMKK2* disruption in CRPC. Two *CAMKK2* knockout (KO) clones were selected (Supplementary Figs. S1A-B) and compared to control (Cas9 only) cells to examine effects on autophagy (Figs. 2D-F). Both *CAMKK2* KO clones exhibited substantially reduced AMPK phosphorylation and LC3B conversion (Fig. 2D) as well as decreased LC3 puncta (Fig. 2E). Compared to control C4-2 Cas9 cells, it was also difficult to find autophagic vesicles in *CAMKK2* knockout cells by TEM (Fig. 2F). However, apoptotic bodies were clearly detectable (Fig. 2F). We confirmed the effects of CAMKK2 inhibition on autophagy using an independent model of CRPC, 22Rv1 cells, in which we created a stable derivative that could express shRNA targeting *CAMKK2* in the presence of DOX. Similar to *CAMKK2* genetic KO in C4-2 cells, the inducible knockdown of CAMKK2 in 22Rv1 cells inhibited autophagy (Figs. 2G-I).

**Fig 2.**
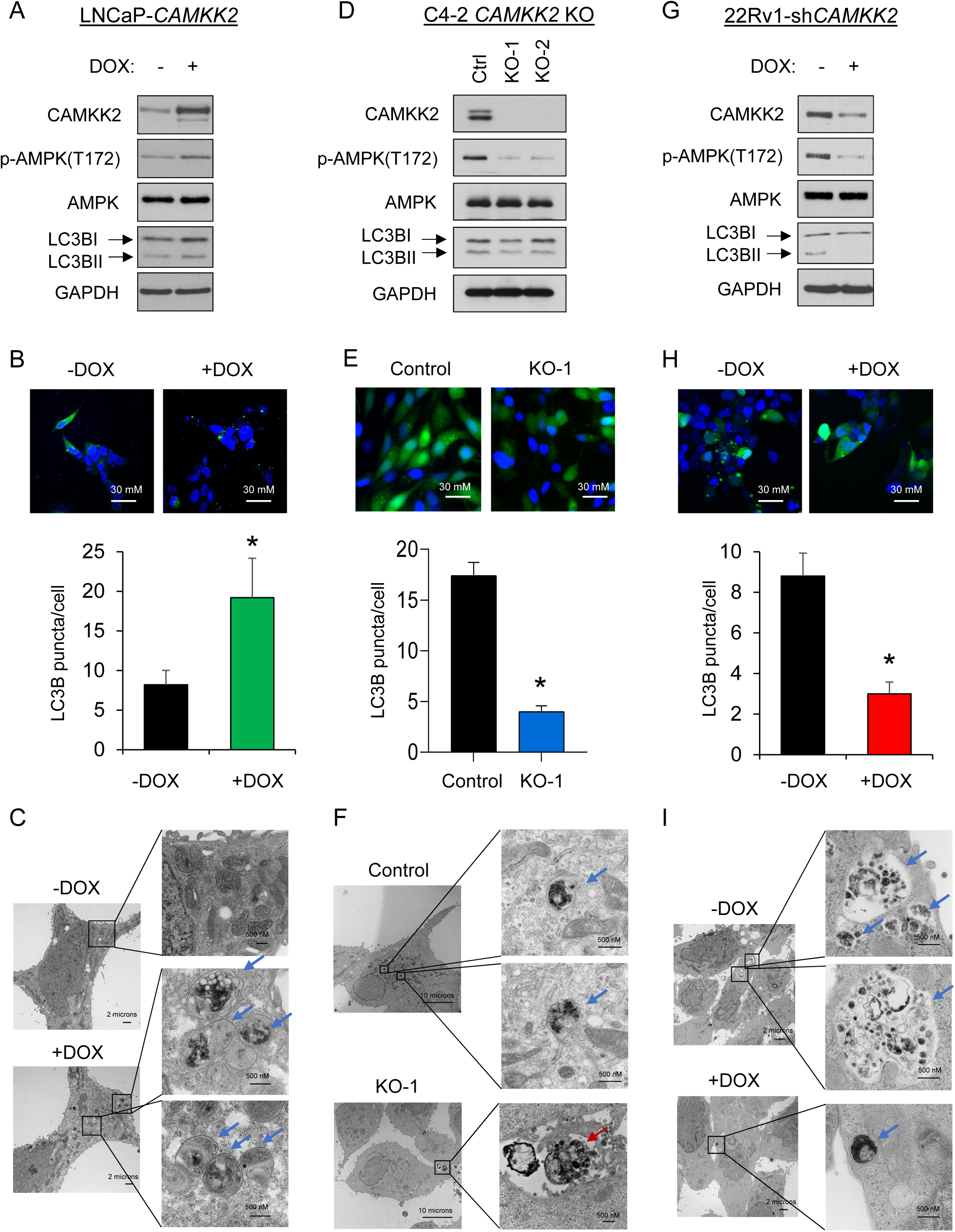
CAMKK2 increases autophagy in prostate cancer cells. (A) Immunoblot analysis of doxycycline (DOX)-inducible LNCaP stable cells (LNCaP-*CAMKK2*) that express CAMKK2 upon addition of 50 ng/ml DOX for 48 hours. (B) LNCaP-*CAMKK2* cells were transiently transfected with GFP-LC3 (green) and then treated ± 50 ng/ml DOX for 48 hours. Representative images (*top*). GFP-LC3 puncta (green) were quantified as the average number of GFP-LC3 puncta per cell ± SEM (*bottom*). The nuclei are stained with DAPI (blue) for reference. *P* value was calculated using a two-tailed *t* test. \**P* < 0.05. (C) LNCaP-*CAMKK2* cells were treated ± 50 ng/ml DOX for 48 hours and imaged using transmission electron microscopy (TEM). Two magnifications of ultrastructures are shown. Blue arrows indicate autophagosomes and autolysosomes. (D) Immunoblot analysis of two independent clones of CRISPR-modified C4-2 *CAMKK2* knockout (KO) cells compared with their parental C4-2 Cas9 control cells (Ctrl). (E) GFP-LC3 was expressed in C4-2 Cas9 control and *CAMKK2* KO cell derivatives. GFP-LC3 puncta (representative images; *top*) and quantification (*bottom*) are shown as in B. (F) C4-2 control and C4-2 *CAMKK2* KO cells were imaged using TEM as in C. Red arrows indicate apoptotic bodies. (G) Immunoblot analysis of DOX-inducible 22Rv1 stable cells that express shRNA targeting *CAMKK2* (22Rv1-sh*CAMKK2*) with 800 ng/ml DOX treatment for 72 hours. (H) 22Rv1-sh*CAMKK2* cells were transiently transfected with GFP-LC3 and then treated ± 800 ng/ml DOX for 72 hours. GFP-LC3 puncta (representative images; *top*) and quantification (*bottom*) are shown as in B. (I) 22Rv1-sh*CAMKK2* cells were treated ± 800 ng/ml DOX for 72 hours and imaged with TEM as in C.

**Supplementary Figure S1**. (A) sgRNAs targeting *CAMKK2* were expressed in C4-2 inducible Cas9 cells and single cell clones were selected after DOX (200 ng/ml) treatment. Immunoblot analysis of these clones and their parental C4-2 Cas9 cells. Note, the clone numbers here were matched for clarity with the numbering throughout the main text and figures. (B) Sanger sequence of C4-2 Cas9 cells and its derivatives *CAMKK2* KO clone 1 and clone 2. Red dashes and letters indicate the identified mutations.

There are several sequential steps involved in autophagy, including initiation, autophagosome formation, autolysosome fusion and degradation. Hence, CAMKK2-mediated increases in LC3 lipidation and relocalization could result from either increased autophagic entry or decreased autophagic flux(25). We therefore used a tandem mCherry-GFP-LC3B fusion protein to evaluate CAMKK2’s role in autophagic flux. LC3B fusion protein is represented as a yellow signal due to the equal expression of both mCherry and GFP basally (diffuse signal) and during early autophagy/prelysosomal fusion (puncta). However, after lysosomal fusion (late autophagy), the acidic environment of the lysosome quenches the GFP signal but retains mCherry, resulting in the colorimetric shift from yellow to red. Consistent with our previous studies(25, 26), androgens increased overall LC3B puncta and GFP^−^mCherry^+^ LC3B puncta (red) (Fig. 3A). We also observed that *CAMKK2* overexpression has a similar result as androgen treatment, which significantly elevated total and red puncta (Fig. 3A). This indicates that CAMKK2, an AR target, can promote autophagic flux similar to androgen treatment. To further validate these findings, we used a lysosomal block assay(25). As described above, chloroquine is a lysosomotropic agent that can block the lysosomal turnover of LC3B. Therefore, impairment of autophagic flux would decrease or not alter LC3BII accumulation in the presence of chloroquine. In contrast, we observed androgens or CAMKK2 expression further increased LC3BII levels in the presence of a lysosomal block, while knockdown of *CAMKK2* decreased LC3B conversion (Fig. 3B and Supplementary Fig. S2), suggesting that CAMKK2 enhanced autophagic flux by increasing autophagy initiation.

**Fig 3.**
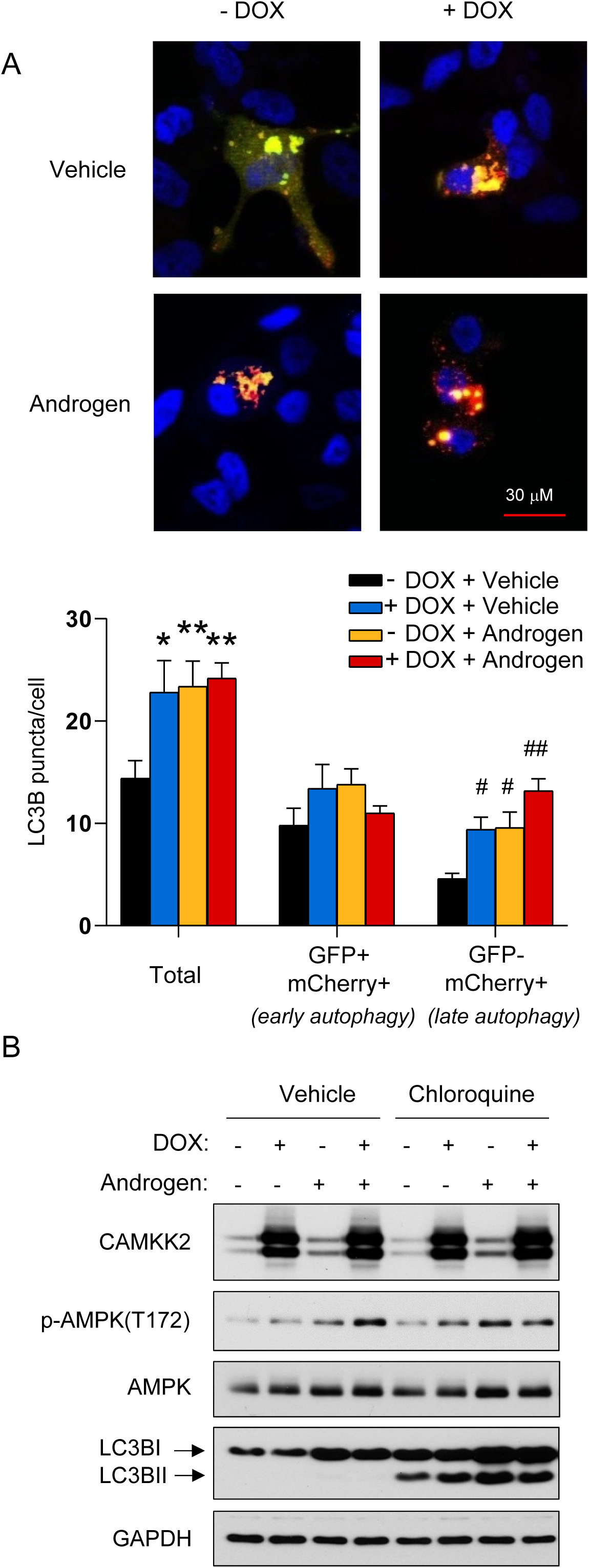
CAMKK2 promotes autophagic flux. (A) LNCaP-*CAMKK2* cells were transfected with an mCherry-GFP-LC3 plasmid and treated ± 10 nM R1881 (androgen) ± 50 ng/ml DOX. Representative fluorescence images of the cellular localization of autophagic puncta (*top*) and quantification (*bottom*). *P* values were calculated using one-way ANOVA with Dunnett’s test. \**P* < 0.05, \*\**P* < 0.01, compared to vehicle group in total. ^#^*P* < 0.05, ^##^*P* < 0.01, compared to vehicle group in GFP-mCherry+. (B) LNCaP-*CAMKK2* cells were treated ± 10 nM R1881 (androgen) ± 50 ng/ml DOX ± 20 μM chloroquine (lysosomal block) for 72 hours. Cell lysates were then subjected to immunoblot analysis.

**Figure S2.** Immunoblot analysis of 22Rv1-sh*CAMKK2* cells ± 800 ng/ml DOX ± 20 μM chloroquine treatment for 72 hours.

### CAMKK2 is required for CRPC cell growth *in vivo*

Previous studies using the pharmacological inhibitor STO-609 have suggested the potential role of CAMKK2 in CRPC growth(4). However, STO-609 has multiple kinase targets(43-45). Here, we used a genetic approach to assess the role of CAMKK2 in CRPC tumorigenesis and progression *in vivo*. Castrated NSG mice were subcutaneously injected with C4-2 Cas9 control and C4-2 *CAMKK2* KO cells (Fig. 4A). *CAMKK2* ablation had a profound effect on CRPC tumor growth (Fig. 4B). In fact, when the average tumor size of the control group was ∼500 mm^3^, no tumors could even be detected in the KO groups. Accordingly, *CAMKK2* KO also dramatically prolonged survival (Fig. 4C). Immunohistochemical (IHC) analysis of tumor tissues determined both a reduction in proliferation and increase in apoptosis in *CAMKK2* KO groups compared to control (Fig. 4D). To validate our findings in a second model of CRPC and test what would happen if we decreased CAMKK2 after tumor implantation, we leveraged our DOX-inducible 22Rv1-sh*CAMKK2* cell model (Fig. 4E and Supplementary Fig. S3A-B). Consistent with the C4-2 *CAMKK2* KO xenograft results, knockdown of *CAMKK2* in 22Rv1 tumors decreased tumor burden over time and consequently increased overall survival (Figs. 4F-G). Moreover, *CAMKK2* knockdown-mediated tumor growth reduction was again correlated to lower proliferation (BrdU) and more apoptosis (TUNEL) (Fig. 4H). Consistent with a pro-survival role of CAMKK2-mediated autophagy, we also observed inducible *CAMKK2* knockdown tumors displayed increased necrosis, but clear regions of perivascular tumor sparing (Fig. 4H, +DOX H&E (high magnification)). Collectively, these data suggest that CAMKK2 is required for maximum CRPC tumorigenesis and progression *in vivo*, potentially by enabling cells to withstand the harsh, nutrient-deficient tumor microenvironment.

**Fig 4.**
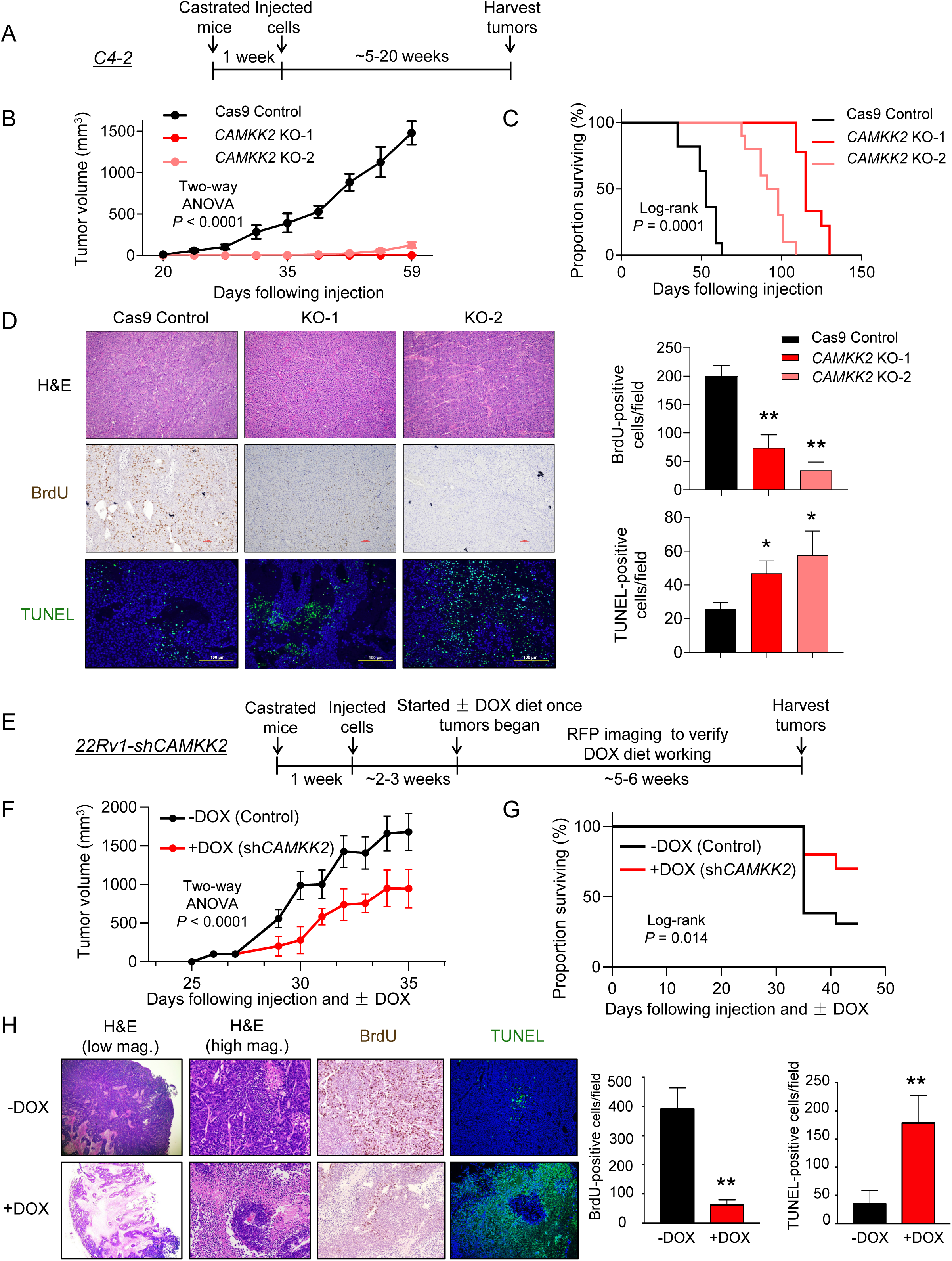
CAMKK2 is required for CRPC tumor growth *in vivo.* (A) Schematic of xenograft study using CRPC C4-2 Cas9 control and *CAMKK2* CRISPR knockout (KO) cell derivatives in castrated NSG mice. (B) Tumor growth curves of C4-2 Cas9 control and C4-2 *CAMKK2* KO xenografts in castrated NSG mice (n = 10/group). *P* values were calculated using two-way ANOVA. (C) Kaplan-Meier survival curve of C4-2 Cas9 control and C4-2 *CAMKK2* KO xenograft mice. *P* values were calculated using the log-rank test. (D) C4-2 xenograft tumor samples were stained with H&E, BrdU and TUNEL. Representative images (*left*) and quantifications of BrdU and TUNEL staining (*right*). \**P* < 0.05, \*\**P* < 0.01 by one-way ANOVA with Dunnett’s test. (E) Schematic of xenograft study using DOX-inducible CRPC 22Rv1-sh*CAMKK2* cells in castrated NSG mice. (F) Tumor growth curves of 22Rv1-sh*CAMKK2* xenografts in castrated NSG mice fed control or DOX-enriched (625 mg/kg) chow. *P* value was calculated using two-way ANOVA. (G) Kaplan-Meier survival curve of 22Rv1-sh*CAMKK2* xenograft mice ± DOX. *P* value was calculated using the log-rank test. (H) 22Rv1-sh*CAMKK2* xenograft tumor samples were stained with H&E, BrdU and TUNEL. Representative images (*left*) and quantifications of BrdU and TUNEL staining (*right*). Note, evidence of perivascular tumor sparing in DOX-treated tumors (H&E high magnification (mag.)). \*\**P* < 0.01 by *t* test.

**Figure S3**. (A) Fluorescence imaging of 5 sample mice (3 on normal chow, 2 on DOX-containing chow) confirming the *CAMKK2* shRNA expression. (B) Immunoblot analysis of tumors from 22Rv1-sh*CAMKK2* xenograft mice ± DOX.

### AR-CAMKK2-AMPK signaling enhanced autophagy through phosphorylation of ULK1 at serine 555

Since AR-CAMKK2 signaling promotes autophagic flux, we further explored the mechanism by which it initiated autophagy. A key protein involved in autophagy initiation is the serine/threonine protein kinase Unc-51 like autophagy activating kinase 1 (ULK1), which functions as part of a complex to transduce upstream signals to the downstream core autophagy machinery(46). AMPK is a known ULK1 upstream regulator by phosphorylating and activating ULK1 at multiple sites in a context dependent-manner(47-49). Thus, we speculated that AR-CAMKK2 activated autophagy through ULK1 in prostate cancer. To explore this possibility, we co-treated LNCaP cells with androgens and the CAMKK2 inhibitor STO-609. Androgens increased the levels of CAMKK2, p-AMPK and LC3BII, an effect that could be abrogated by STO-609 (Fig. 5A). Androgens also increased ULK1 phosphorylation at serine 555, an effect that was again reversed by STO-609 (Fig. 5A). This was of interest because serine 555 has been shown to be a critical phosphorylation site necessary for AMPK-mediated autophagy *in vitro* and *in vivo* (47, 50-52). To exclude the non-specific effects of STO-609, we next tested p-ULK1(S555) status in cells following genetic or molecular modification of CAMKK2 and AMPK. In LNCaP-*CAMKK2* cells, DOX alone increased CAMKK2 expression level, resulting in a similar increase in p-AMPK, p-ULK1 and LC3BII levels compared to androgen treatment alone (Fig. 5B). These increases could be reversed upon knockdown of AMPKα1, the predominant AMPK α catalytic subunit in prostate cancer(3, 9, 53, 54). The requirement for AMPK was confirmed with three independent siRNAs (Fig. 5C). To verify that ULK1 phosphorylation was necessary for CAMKK2-mediated autophagy, we transfected LNCaP-*CAMKK2* cells with constructs expressing vector control, WT ULK1 or ULK1 4SA mutant, an AMPK non-phosphorylatable ULK1(47). Cells treated with DOX (CAMKK2 expression) and expressing WT ULK1 had increased LC3BII levels, indicating an increase of autophagy, while 4SA mutant blocked CAMKK2-mediated autophagy (Fig. 5D). The requirement of CAMKK2 for AMPK-ULK1-mediated autophagy was confirmed in both the C4-2 and 22Rv1 CRPC models (Figs. 5E-F). Taken together, these findings demonstrated that AR-CAMKK2 triggers AMPK phosphorylation and activation, and in turn phosphorylates ULK1 at serine 555, which ultimately stimulates autophagy.

**Fig 5.**
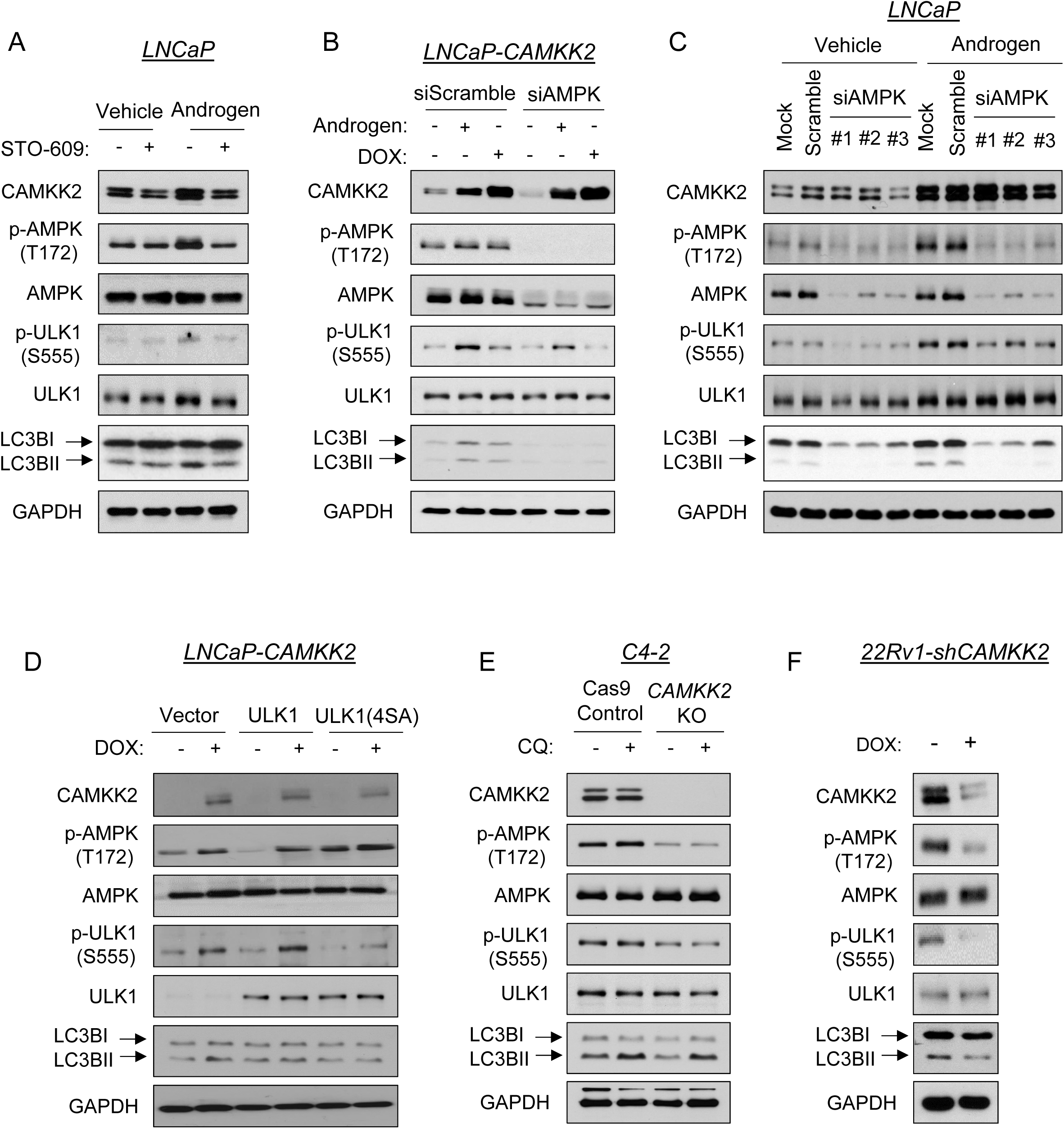
AR-CAMKK2-AMPK signaling increases autophagy by phosphorylating ULK1 at serine 555. (A) LNCaP cells were treated ± 10 nM R1881 (androgen) ± 30 μM STO-609 for 72 hours. (B) LNCaP-*CAMKK2* cells were transfected with siRNAs targeting scramble control or the α1 catalytic subunit of AMPK (si*AMPK*) and then treated with androgen for 72 hours or DOX (50 ng/ml) for 48 hours. Cell lysates were subjected to immunoblot analysis. (C) Parental LNCaP cells were transfected with siRNAs targeting scramble control or three different regions of the α1 catalytic subunit of AMPK (si*AMPK*) and then treated with vehicle or androgen for 72 hours. Cell lysates were subjected to immunoblot analysis. (D) LNCaP-*CAMKK2* cells were transfected with empty vector, ULK1 or ULK1 (4SA) expression constructs and then treated ± DOX for 48 hours. Cell lysates were subjected to immunoblot analysis. (E) Immunoblot analysis of C4-2 Cas9 control and *CAMKK2* KO derivative cells treated with vehicle or chloroquine (20 μM). (F) Immunoblot analysis of 22Rv1-sh*CAMKK2* cells ± 800 ng/ml DOX treatment for 72 hours.

### ULK1 correlates with poor patient prognosis in men with prostate cancer

To examine the clinical association between ULK1 and patient prognosis, we analyzed two well-annotated, publicly available patient databases. The expression level of *ULK1* mRNA was inversely correlated with disease-free survival in both TCGA(55) (Fig. 6A) and Taylor *et al*. 2010(56) (Fig. 6B) clinical cohorts. Consistent with these clinical correlations, previous histological studies have linked high ULK1 levels to biochemical recurrence and PSA levels(57, 58). Taken together, these data suggest that high ULK1 may promote disease progression.

**Fig 6.**
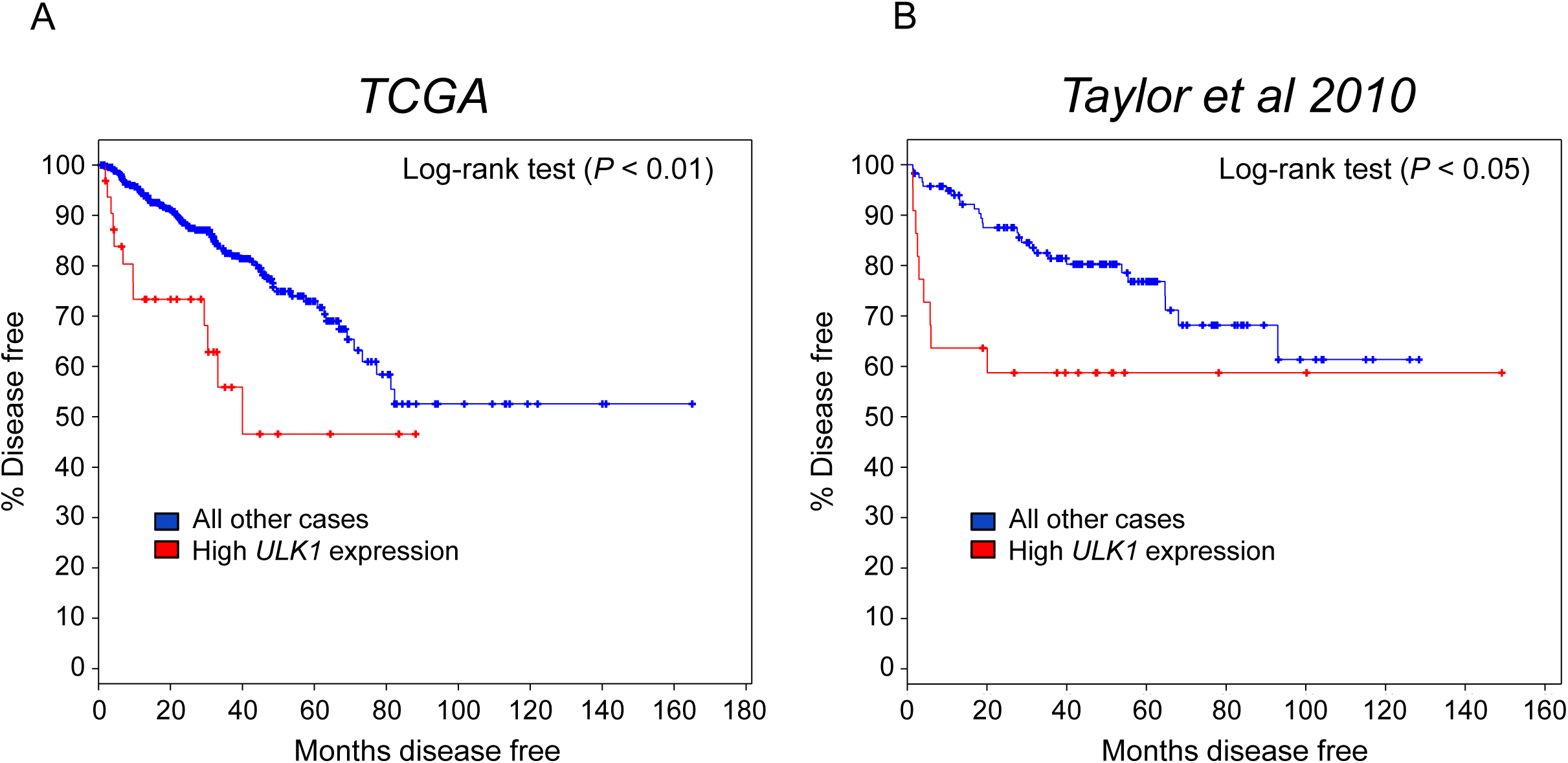
High *ULK1* tumor expression predicts poor patient prognosis in independent clinical cohorts of men with prostate cancer. Kaplan-Meier estimates of disease-free survival in The Cancer Genome Atlas (TCGA) and Taylor *et al.* 2010 clinical cohorts based on *ULK1* expression. Data were generated from cBioPortal.

### Pharmacological targeting of ULK1 inhibits prostate cancer cell growth

We next wanted to determine if ULK1 was a potential therapeutic target in prostate cancer. To test this, we leveraged SBI-0206965 (6965), a recently described ULK1 inhibitor that has shown anti-cancer effects in lung cancer cells under nutrient deprivation (Fig. 7A)(59). To validate 6965’s antagonistic effects in prostate cancer cells, we first used the known ULK1 substrate VPS34 to determine whether 6965 could block ULK1 activity. 6965 decreased both basal and androgen-induced phosphorylation of VPS34 at serine 249, in alignment with the reduction of LC3BII (Fig. 7B). In 22Rv1 cells, 6965 also resulted in inhibition of p-VPS34 and LC3BII accumulation (Supplementary Fig. S4). Interestingly, we noticed an increase of p-AMPK after 6965 treatment, consistent with the previously described negative feedback loop that exists between ULK1 and AMPK(60). Next, to assess the efficacy of 6965 on prostate cancer cell growth, we treated LNCaP-*CAMKK2* cells with androgens, DOX and/or 6965 for 7 days. Although 6965 did not significantly inhibit basal LNCaP cell growth, it blocked androgen- and/or DOX-mediated LNCaP cell growth (Fig. 7C), consistent with our previous findings that siRNA-mediated knockdown of ULK1 blocked androgen-mediated cell growth(26). Interestingly, the LNCaP-derived CRPC derivative C4-2 cells were more sensitive to 6965 treatment, showing a ∼70% reduction in growth (Fig. 7D). In 22Rv1 cells, ∼50% growth inhibition was observed (Fig. 7E). To evaluate the long-term effects of 6965 on cell proliferation, we performed clonogenic assays. All three cells were very sensitive to prolonged 6965 treatment with almost 100% inhibition in clonogenic potential (Figs. 7F-H). Collectively, these data indicate that ULK1 is a potentially druggable target for the treatment of prostate cancer. Future studies would need to explore the safety of such an approach *in vivo*.

**Fig 7.**
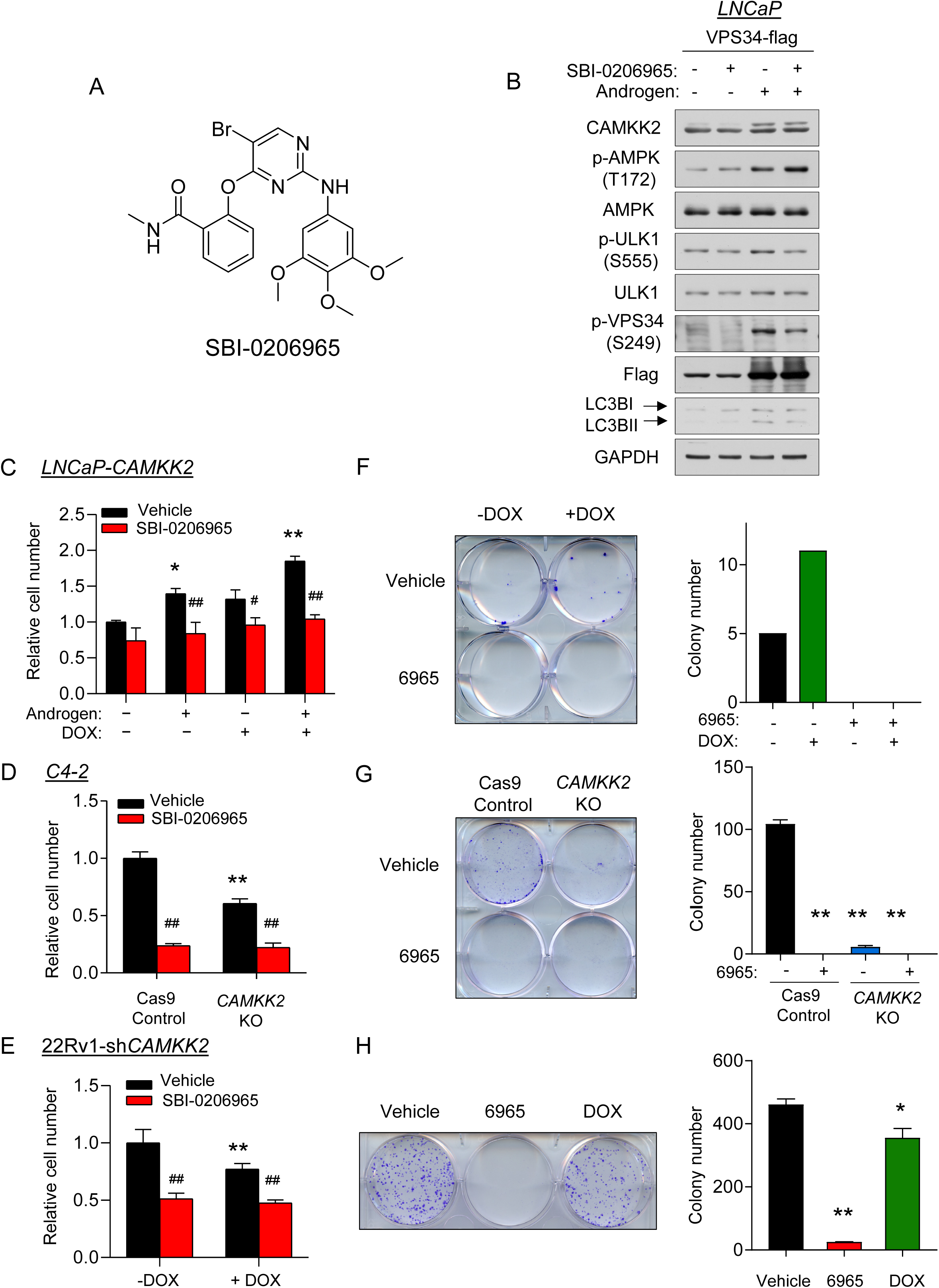
The ULK1 inhibitor SBI-0206965 represses prostate cancer cell growth. (A) Chemical structure of the ULK1 inhibitor SBI-0206965. (B) LNCaP cells were transfected with VPS34-FLAG following 72 hours 10 nM R1881 (androgen) treatment. Cell lysates were collected 2 hours after vehicle or SBI-0206965 (10 μM) treatment and subjected to immunoblot analysis. (C) Cell growth of LNCaP-*CAMKK2* cells following 7 days R1881 (androgen, 10 nM), DOX (50 ng/ml) and/or SBI-0206965 (10 μM) treatment. \**P* < 0.05, \*\**P* < 0.01 compared to no androgen/DOX/SBI-0206965 treatment group. ^#^*P* < 0.05, ^##^*P* < 0.01, compared to corresponding vehicle (SBI-0206965) treatment group. (D) Cell growth of C4-2 Cas9 control and C4-2 *CAMKK2* KO derivative cells ± SBI-0206965 (10 μM). \*\**P* < 0.01 compared to C4-2 control cells. ^##^*P* < 0.01, compared to vehicle treatment group. (E) Cell growth of 22Rv1-sh*CAMKK2* cells treated for 7 days ± DOX (800 ng/ml) ± SBI-0206965 (10 μM). \**P* < 0.05, \*\**P* < 0.01 compared to no DOX treatment group. ^##^*P* < 0.01, compared to corresponding vehicle (SBI-0206965) treatment group. (F) Colony formation assay of LNCaP-*CAMKK2* cells following 28-day DOX and/or SBI-0206965 (10 μM) under 100 pM R1881 (androgen) treatment (required for LNCaP colony formation). Representative image (*left*). Quantification (*right*). (G) Colony formation assay of C4-2 Cas9 control and C4-2 *CAMKK2* KO derivative cells ± SBI-0206965 (10 μM) for 21 days. Representative image (*left*). Quantification of three independent experiments (*right*). \*\**P* < 0.01, compared to C4-2 control vehicle treatment group. (H) Colony formation assay of 22Rv1-sh*CAMKK2* cells treated for 21 days ± DOX (800 ng/ml) or SBI-0206965 (10 μM). Representative image (*left*). Quantification of three independent experiments (*right*). \**P* < 0.05, \*\**P* < 0.01, compared to vehicle treatment group.

**Figure S4**. 22Rv1 cells were transfected ± VPS34-FLAG for 48 hours. Immunoblot analysis of transfected cells with 2 hours SBI-0206965 (10 μM) treatment.

## Discussion

Although autophagy has context-dependent roles in cancer (18, 61-64), our data support a pro-cancer role for this cellular process in prostate cancer. These findings are consistent with our previous work(25, 26) and the work of others in the field(27, 28, 63, 65-67). As presented in our previous reports, blocking autophagy by molecular or pharmacological approaches resulted in decreased androgen-mediated prostate cancer cell growth(25, 26). Mechanistically, androgens stimulate AR to promote autophagy through multiple mechanisms including the indirect accumulation of intracellular ROS and more directly through the transcription of several core autophagy genes(25, 26). In this study, we revealed a novel mechanism underlying how AR regulates autophagy. Our data demonstrated that an AR-CAMKK2-AMPK signaling cascade can drive autophagy through the phosphorylation of ULK1, an important initiator of autophagy, at serine 555. This phosphorylation activates the ULK1 complex and ultimately initiates autophagy and autophagic flux for prostate cancer cell proliferation and survival (Fig. 8). This finding not only provides a novel mechanistic insight into AR’s regulation of autophagy, but highlights potential new avenues for therapeutic targeting of autophagy in prostate cancer. A non-AR-mediated regulation of autophagy has been reported as a resistance mechanism to treatment with the anti-tumor compound triptolide in prostate cancer(68). As a result, chloroquine was applied to overcome triptolide resistance, enhancing the anti-tumor effect of triptolide in much the same way chloroquine enhanced the effect of hormone ablation in our own CRPC models (Fig. 1). Despite differences identified in the ULK1 phosphorylation sites, our results agree with the overall concept that CAMKK2-AMPK-induced ULK1 activation and autophagy provides an important survival mechanism for prostate cancer cell growth.

**Fig 8.**
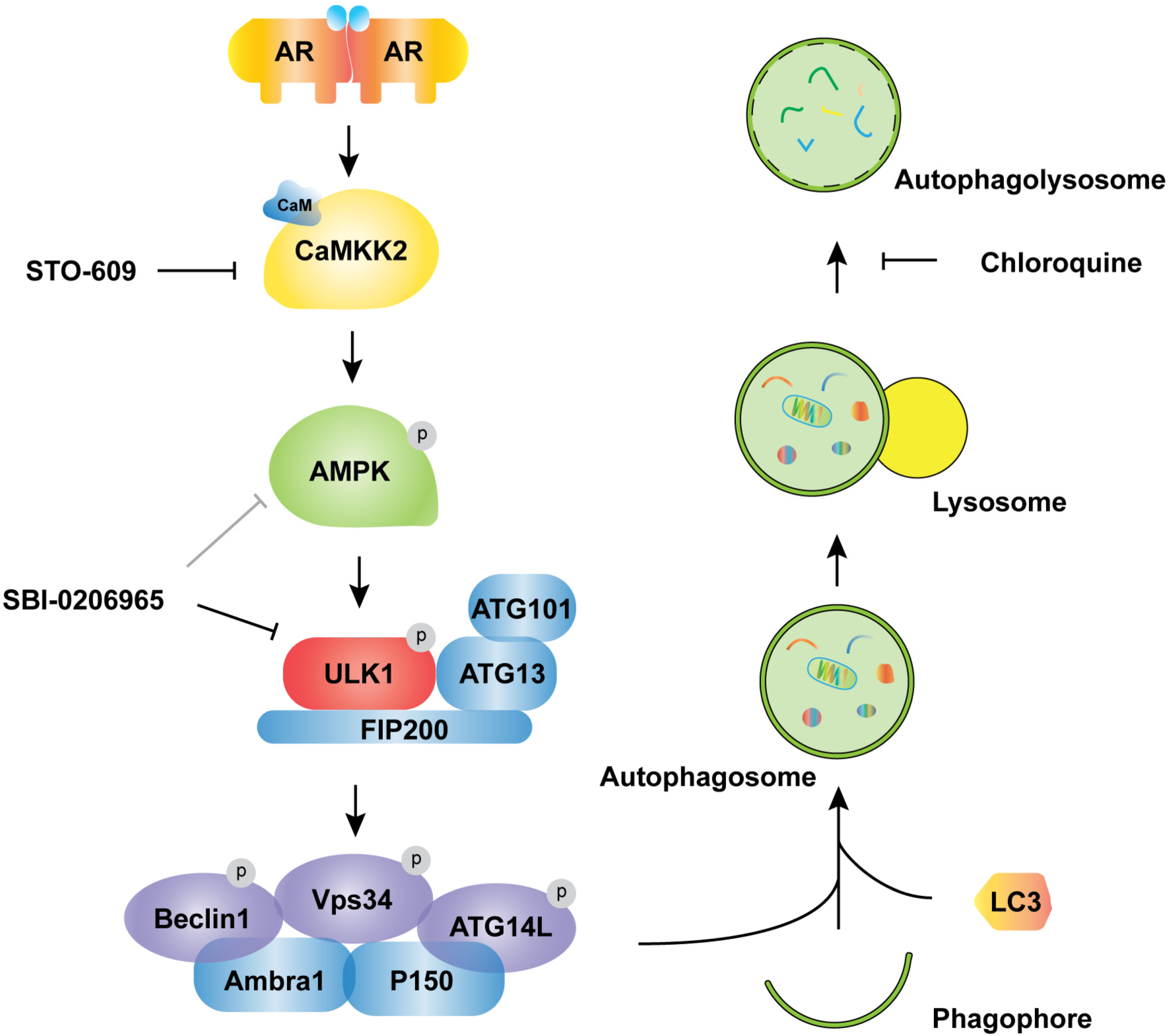
Working model depicting how AR-CAMKK2-AMPK signaling regulates autophagy by ULK1 phosphorylation and activation in prostate cancer. AR increases the expression of *CAMKK2* which in turn phosphorylates and activates AMPK at threonine 172. As a result, AMPK phosphorylates ULK1 at serine 555 which activates the ULK1 complex and initiates autophagy, supporting prostate cancer growth. This growth and survival mechanism can be blocked at several steps and as such, offers alternative strategies for targeting autophagy in prostate cancer.

Interestingly, despite agreement that autophagy promotes prostate cancer progression, how this process is regulated by AR is still debated(18, 25, 26, 28, 36, 53, 63, 67, 69, 70). These discrepancies may be attributable to differences in the duration of upstream signals, reliance on indirect or nonselective modulators of autophagy or treatment conditions. As we and others have shown, androgens, in an AR-dependent mechanism, can directly and indirectly increase autophagy through a variety of mechanisms including elevating intracellular ROS levels and transcription of core autophagy genes(25-27, 63). As shown here, there is also a clear, direct AR regulation of AMPK-mediated autophagy through the expression of *CAMKK2*. The mechanism underlying how antiandrogens can, like androgens, paradoxically also can increase autophagy is less clear. But these different observations may speak to the potential benefit of targeting downstream effector processes like autophagy that can be activated under a variety of conditions to drive disease progression. Our data presented here provides evidence that targeting CAMKK2-AMPK-ULK1 signaling may be an effective, alternative strategy to block protective autophagy in advanced prostate cancer.

Under glucose or amino acid starvation, ULK1 is well characterized to be regulated by AMPK. AMPK binds to the serine/proline-rich domain and can phosphorylate ULK1 at multiple sites (S317, S467, S555, T575, S637 and S777) which subsequently change ULK1 conformation and enhance its kinase activity. This, in turn, promotes the formation of the ULK1 complex (ULK1, ATG13, ATG101, and FIP200)(41, 46). Activated ULK1 can further phosphorylate downstream VPS34 complex members to induce autophagic entry(46). In this study, we first demonstrated the S555 site of ULK1 as a downstream target of AMPK in response to androgen treatment. S555 was increased under androgen treatment but could not be activated when cells were subjected to *AMPK* siRNA (Fig. 5B&C). When cells were reconstituted with a non-phosphorylatable ULK1 mutant (4SA), they were defective in autophagy following AMPK activation (Fig. 5D). Although we cannot exclude contributions from other phosphorylation sites, these findings suggest the functional importance of ULK1 S555 by AMPK in AR-mediated autophagy induction and support prior reports that S555 is functionally one of the most important AMPK target sites on ULK1(47, 51, 52).

Interestingly, we observed a negative feedback loop between AMPK and ULK1 similar to what has been described before in HEK293 cells under starvation(60). While non-phosphorylatable ULK1 mutants impaired autophagy, they significantly increased p-AMPK (T172) (Fig. 5D). Likewise, when cells were treated with the ULK1 inhibitor SBI-0206965, a robust enhancement of p-AMPK was detected (Figs. 7B and Supplementary Fig. S4). It is unclear at this time if this translates to other AMPK-mediated processes being hyperactivated and therefore influencing prostate cancer cell pathobiology.

The efficacy of autophagy inhibition in preclinical models of cancer has paved the way for new clinical trials investigating the efficacy of autophagy inhibition in patients, particularly in combination with traditional anti-cancer treatments. Chloroquine and its derivative hydroxychloroquine, as FDA-approved drugs, have been favored and repurposed in prostate cancer. Previous studies indicated that chloroquine in combination with other therapeutic agents including anti-androgens, chemotherapy and kinase inhibitors can induce greater cytotoxicity than single agent treatment alone(15, 67, 69, 71-73). Likewise, our data indicate anti-cancer effects for chloroquine in combination with androgen deprivation therapy (Fig. 1). Although a series of clinical trials in prostate cancer have been started to test the efficacy of chloroquine analogs, thus far, limited clinical efficacy has been observed. This is believed to be due in large part to an inability to achieve the drug concentration needed for sustained inhibition of autophagy within tumors prior to the onset of significant side effects(16). Despite the fact that hydroxychloroquine is safer than chloroquine, a micromolar concentrations are required to maintain autophagy inhibition in patients(71). Even so, variable effects on autophagy are still being observed, possibly due to inconsistencies in cell penetration that are in part dependent on the individual’s tumor microenvironment(15). Thus, long-term and high-dosage treatments will inevitably reduce the therapeutic window. Given the potential challenges in the use of lysosomotropic agents, which are not even specific for autophagy, targeting other steps in autophagy, such as ULK1, may provide alternative solutions.

ULK1 expression is highly correlated with patient disease-free time, biochemical recurrence, Gleason score, and metastasis (Fig. 6 &(57, 58)). Currently, three studies have investigated this ULK1 inhibitor and showed selective and potent inhibition on ULK1 activity(59, 74, 75). In agreement with other reports, our data showed that SBI-0206965 inhibited ULK1 activity as evidenced by the reduction of p-VPS34 (S249) (Figs. 7B and Supplementary Fig. S4). Moreover, SBI-0206965 exhibited its anti-growth activity in both hormone-sensitive and CRPC cells (Figs. 7C-H). A recent study suggested that SBI-0206965 is a dual inhibitor of AMPK and ULK1(76). While this would potentially be beneficial for blocking two important nodes of AR-CAMKK2-AMPK-ULK1 signaling, we did not observe a consistent decrease of p-ULK1 after SBI-0206965 treatment, suggesting SBI-0206965 may not function as an AMPK inhibitor in our models. However, we acknowledge that this interpretation may be convoluted due to the above-described feedback mechanism between AMPK and ULK1(60). In addition, the efficacy and pharmacokinetic profile of SBI-0206965 *in vivo* are still largely unknown.

Given that systemic blocking or genetic ablation of CAMKK2 appears well-tolerated in mouse models and CAMKK2 has a more restricted expression profile but is elevated in prostate cancer, we propose targeting CAMKK2 may be a viable alternative. Unfortunately, the use of STO-609 as used in this study is likely not a clinically viable option due to its off-target effects on other kinases and pharmacokinetic limitations(43-45). There are, however, ongoing efforts to develop next-generation CAMKK2 inhibitors(44, 77-79).

In summary, our results provide a novel mechanism that links AR signaling and protective autophagy in prostate cancer. Targeting CAMKK2 decreases the AMPK-mediated phosphorylation of ULK1 at serine 555, which in turn stalls the initiation of autophagy and impairs prostate cancer cell growth. These findings not only add a mechanistic layer of complexity to shed light on AR’s regulation of autophagy, but also provides new opportunities for inhibiting autophagy in prostate cancer that we postulate warrant being tested to determine if they can overcome the existing limitations of chloroquine.

## Materials and methods

### Cell culture, plasmids and reagents

LNCaP, 22Rv1 and HEK293T cell lines were obtained from American Type Culture Collection (Baltimore, MD, USA) (CRL-1740, CRL-2505, CRL-3216). C4-2 cells were obtained from Dr. Nancy Weigel (Baylor College of Medicine). LNCaP-*CAMKK2* and 22Rv1-sh*CAMKK2* cells have previously been described(8). 22Rv1-fLuc cells were created by pBABE-fLuc-YFP plasmid (a gift from Dr. Christopher Counter, Duke School of Medicine) with retroviral transduction strategy(80). Cells were maintained as previously described(26) and validated by STR profiling (University of Texas MD Anderson Cancer Center Cell Culture Core). All cells were confirmed to be mycoplasma-free by MycoAlert Mycoplasma Detection Kit (Lonza, Morristown, NJ USA; Cat #: LT07-118). Cells were steroid-starved in phenol red-free medium containing 10% charcoal stripped-FBS (5% CS-FBS for C4-2 cells) for 72 hours before treatment unless otherwise noted. pCW-Cas9 and pLX-sgRNA were gifts from Drs. Eric Lander & David Sabatini (Addgene, Watertown, MA, USA; plasmids #: 50661, 50662). pcDNA4-VPS34-Flag was a gift from Dr. Qing Zhong (Addgene plasmid #: 24398). pcDNA3.1-hULK1 and 4SA mutant were gifts from Dr. Mondira Kundu (St. Jude Children’s Research Hospital). Enhanced GFP-LC3 and mCherry-GFP-LC3B constructs have been previously described(81). The synthetic androgen methyltrienolone (R1881) was purchased from PerkinElmer (Naperville, IL, USA; Cat #: NLP005005MG). Chloroquine (Cat #: C6628), doxycycline hyclate (Cat #: D9891), puromycin (Cat #: P8833), BrdU (5-bromo-2-deoxyuridine, Cat #: B5002) and polybrene (Cat #: TR-1003) were purchased from Sigma-Aldrich (St. Louis, MO, USA). G418 sulfate was purchased from Gold Biotechnology (St. Louis, MO, USA; Cat #: G-418-25). Blasticidin was purchased from Millipore Sigma (St. Louis, MO, USA; Cat #: 203350).

### Xenografts, histology and immunostaining

All animal experiments were approved by and conducted under the Institutional Animal Care and Use Committee (IACUC) at the University of Texas MD Anderson Cancer Center and the University of Houston according to NIH and institutional guidelines. Xenografts were performed on 6-8 weeks male NSG mice obtained from either The Jackson Laboratory (Bar Harbor, ME, USA; Cat #: 005557; Fig. 1&3E) or The University of Texas MD Anderson Cancer Center Experimental Radiation Oncology Breeding Core (Fig. 3A). Castrations were conducted one week before injections. One million cells in 200 μl DPBS: Matrigel® 1:1 vol/vol (Corning, Corning, NY, USA; Cat #356231) were injected subcutaneously into flanks. Tumor size was measured by calipers until tumor lengths in the control group reached 1.5 cm or signs of morbidity were observed (ex. reduced body weight or hunched back). Tumor volume was calculated by the formula: length x width^2^/2.

For 22Rv1-sh*CAMKK2* xenografts, mice were randomized into normal/control or doxycycline-containing (625 mg/kg, Envigo, IN, USA) diet groups. Then, shRNA expression with surrogate red fluorescent protein (RFP) was tracked by fluorescence (IVIS Spectrum *In Vivo* Imaging Station, PerkinElmer). For chloroquine xenograft experiments, mice were randomly grouped into vehicle control or chloroquine IP treatment when the tumor volume reached 100 mm^3^. One hour before tissue/tumor collection/sacrifice, mice were injected with 100 mg/kg BrdU. Half of the tumor sample was snap frozen while the other half was immediately fixed in 4% PFA overnight at 4°C. For staining, samples were dehydrated and embedded in paraffin. Paraffin slides were then rehydrated and further processed with antigen retrieval in citrate buffer (DAKO, Santa Clara, CA, USA; Cat #: S169984-2). Peroxidase blocking was performed in 1% H_2_O_2_ plus 10% methanol solution. Proliferative cells were detected by BrdU staining. For this, slides were blocked with goat serum (DAKO; Cat #: X090710-8) and incubated overnight with anti-BrdU antibody (Calbiochem: Part of Millipore Sigma; Cat #: NA61). After washing with PBST (PBS with 0.02% Tween 20), secondary antibodies (Mouse-on-Mouse HRP Polymer, Biocare Medical, CA, USA, Cat#: MM620) were incubated for 30 minutes. Sections were developed by DAB (Vectorlabs, Burlingame, CA, USA; Cat #: SK-4100). Apoptotic cells were detected by TUNEL staining using the *In Situ* Cell Death Detection Kit, Fluorescein (Roche, Madison, WI, USA; Cat #: 11684795910) following the manufacturer’s instructions. Hematoxylin and eosin staining was performed by the University of Texas MD Anderson Cancer Center Department of Veterinary Medicine and Surgery Research Animal Support Facility. Microscopy was done with an Olympus BX51 microscope and cellSens imaging software (Olympus, Center Valley, PA, USA). Analysis was done on 3-6 acquired fields per section and data were averaged.

### Plasmid and small interfering RNA (siRNA) transfections

All transfections were conducted as previously described(8, 81). In brief, plasmids were transfected using Lipofectamine 2000 transfection reagent (Thermo Fisher Scientific, Waltham, PA, USA) according to the manufacturer’s instructions. The siRNAs were purchased from Thermo Fisher Scientific and transfected using DharmaFECT 1 transfection reagent. Sequences of the shRNAs and siRNAs used in this study are listed in Supplementary Table 1.

### Generation of CRISPR/Cas9 *CAMKK2* knockout cells

pCW-Cas9 was co-transfected with lentiviral packaging plasmids into actively growing HEK293T cells using Lipofectamine 2000 transfection reagent. After 48 hours, medium containing virus was collected, filtered and added to the target cells with 8 μg/ml polybrene. After 48 hours, fresh medium with 1 μg/ml puromycin was used to select doxycycline-inducible Cas9 expressed target cells. The gRNAs targeting *CAMKK2* were designed by http://crispor.tefor.net/(82) and synthesized by Sigma (listed in Supplementary Table 1). The sgRNA oligos were cloned into pLX-sgRNA. pLX-*CAMKK2* sgRNAs were transfected into Cas9-inducible expressing cells by the same lentivirial transduction strategy before selection with 10 μg/ml blasticidin. Cells expressing inducible Cas9 and sgRNA were first treated with doxycycline for 7 days. This method limited the Cas9 activation window and therefore greater potential for off-target CRISPR effects. After, single clones were isolated and screened to establish *CAMKK2* knockout cells. Parental Cas9-inducible cells were used as control. Each clone was validated by sequencing and western blot.

### Western blot analysis

Western blot analysis was performed as previously described(8, 9, 26, 81). Briefly, cells were harvested in RIPA lysis buffer (50 mM Tris HCl pH 8, 150 mM NaCl, 1% NP-40, 0.5% sodium deoxycholate, 0.1% SDS) supplemented with Complete Protease Inhibitor Cocktail (Sigma, Cat #: 11697498001) and PhosSTOP phosphatase inhibitor (Roche, Cat #:4906845001). Primary antibodies were purchased from the following sources: Cell Signaling Technology (Danvers, MA, USA): ULK1 (Cat #: 4773), p-ULK1(S555) (Cat #: 5869), LC3B (Cat #: 2775), p-AMPK(T172) (Cat #: 2535), p-VPS34(S249) (Cat #: 13857); Sigma: CAMKK2 (Cat #: HPA017389), GAPDH (Cat #: G8795), FLAG (Cat #: F1804).

### Immunofluorescence microscopy

GFP-LC3/mCherry-GFP-LC3 fusion constructs were expressed in cells as previously described(26, 81). Following treatments, cells were fixed with 4% PFA for 15 min at RT and DAPI was used as a counterstain. Images were captured using the Olympus BX51 fluorescence microscope and cellSense imaging software. Samples were analyzed by Image J where LC3 puncta per cell were counted for 50 cells per cell line and averaged.

### Transmission electron microscopy (TEM)

Cells were plated at 100,000 cells/well in 6-well plates and treated as indicated in figures/figure legends. Samples were fixed with a Karnovsky’s fixative solution (3% glutaraldehyde plus 2% paraformaldehyde in 0.1 M cacodylate buffer, pH 7.3) at 4 °C. Samples were further processed by the University of Texas MD Anderson Cancer Center High Resolution Electron Microscopy Core Facility.

### Proliferation assays

Proliferation assays were carried out as previously described by measuring the cellular double-stranded DNA content using a fluorescent DNA stain(81).

### Clonogenic assays

Cells were plated at 5000 (LNCaP) or 1000 (C4-2, 22Rv1) cells/well in 6-well plates. Colonies were formed for 3-4 weeks. Media and treatments were refreshed every week. Cells were fixed with acetic acid/methanol 1:7 (vol/vol) and then stained with 0.5% crystal violet. The number of visible colonies were counted. The data were representative of three independent experiments with similar results.

### Statistical analysis

Statistical analyses were performed using Microsoft Excel 2013 (Redmond, WA, USA) and GraphPad Prism 8 (San Diego, CA, USA). Bioinformatic analyses of the correlation of ULK1 gene expression with patient prognosis were generated from cBioPortal(83, 84). One-way or two-way ANOVAs, Student *t*-tests were used to determine the significance among groups where appropriate as indicated in the figures or figure legends. Log-rank test was used to determine the significance of Kaplan-Meier curves. Grouped data are presented as mean α SEM unless otherwise noted. *P* values are indicated in figures or figure legends.

## Acknowledgments

We thank members of the Frigo laboratory for helpful discussions and editorial assistance during the preparation of the manuscript. We also thank Dr. Mondira Kundu (St. Jude Children’s Hospital) for the gift of the ULK expression constructs, Dr. Christopher Counter (Duke School of Medicine) for the gift of pBABE-fLuc-YFP construct, Dr. Nancy Weigel (Baylor College of Medicine) for the C4-2 cell line, Kenneth Dunner, Jr. in the High Resolution Electron Microscopy Core Facility at MD Anderson Cancer Center for his help in acquiring the TEM images, and Dr. Jeff Spencer (University of Houston) for the help with sgRNA design. This work was supported by grants from the National Institutes of Health (NIH R01CA184208 to D.E.F.), American Cancer Society (RSG-16-084-01-TBE to D.E.F.), an Institutional Research Grant (to D.E.F.), startup grants from the University of Texas MD Anderson Cancer Center (to D.E.F.) and generous philanthropic contributions to The University of Texas MD Anderson Moon Shots Program (to D.E.F.). This work was also supported by an Antje Wuelfrath Gee and Harry Gee, Jr. Family Legacy Scholarship (to C.L.) and MD Anderson Odyssey fellow supported by the CFP foundation (to A.B.B). Electron microscopy was performed by the CCSG-funded University of Texas MD Anderson Cancer Center High Resolution Electron Microscopy Facility while histology was performed in conjunction with the CCSG-funded University of Texas MD Anderson Cancer Center Research Histology Core Laboratory, NIH grant P30CA0166772.

