## Supplemental Material for "Inhibition of CAMKK2 impairs autophagy and castration-resistant prostate cancer via suppression of AMPK-ULK1 signaling"

### Figure S1

A

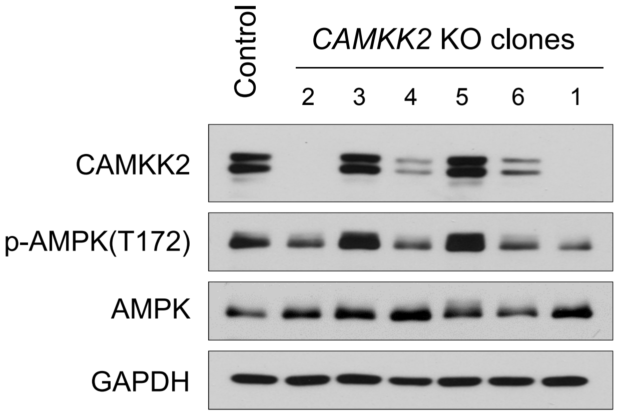


B

**C4-2 Cas9 Control (WT *CAMKK2*)**

5’-CCGTGACATCAAACCTTCCA-3’

**KO clone-1 (Heterozygous)**

5’-CCG—ACATCAAACCTTCCA-3’ (deletion)

5’-CCG<u>G</u>TGACATCAAACCTTCCA-3’ (insertion)

**KO clone-2 (Homozygous)**

5’-CCG<u>T</u>TGACATCAAACCTTCCA -3’ (insertion)

### Figure S2

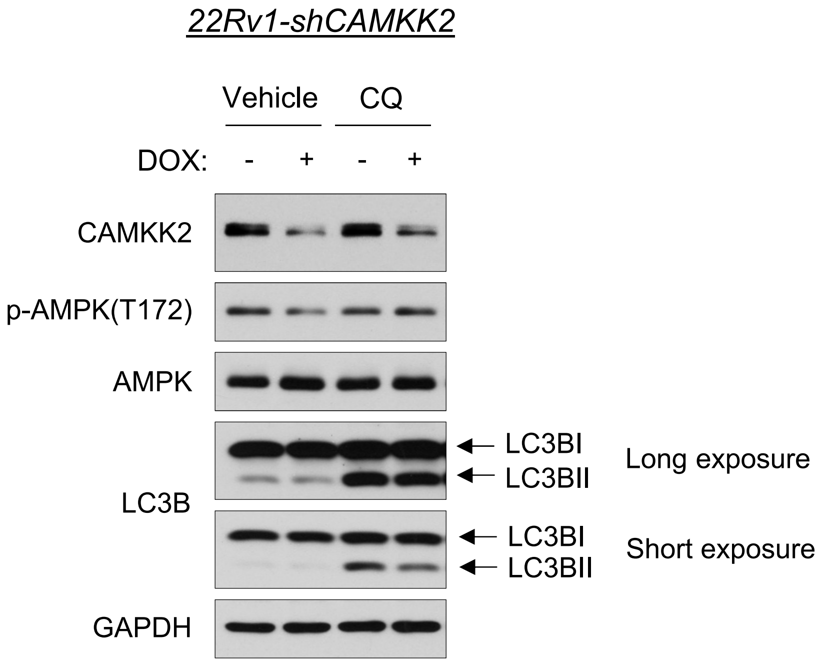
22Rv1-shCAMKK2
Vehicle
CQ
DOX:
-
+
-
+
CAMKK2
p-AMPK(T172)
AMPK
LC3B
LC3BI
LC3BII
Long exposure
LC3BI
LC3BII
Short exposure
GAPDH

### Figure S3

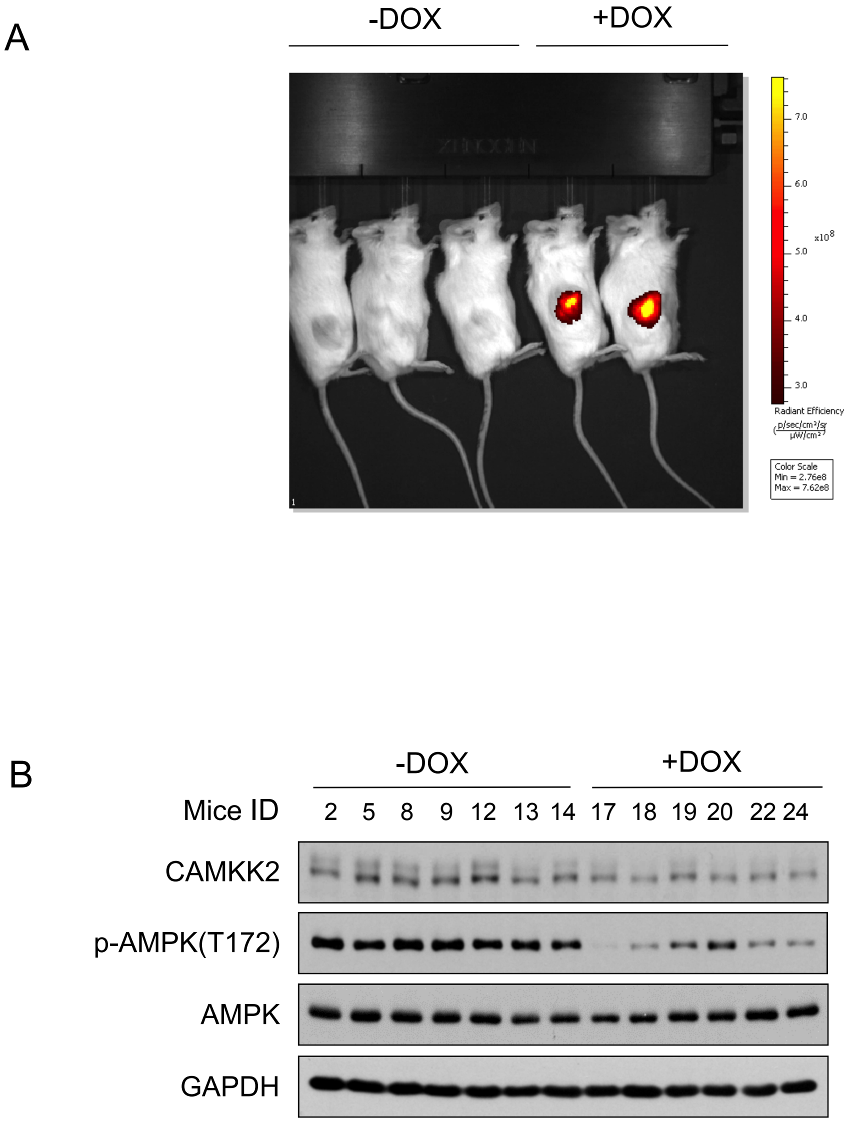
A
-DOX
+DOX
Radiant Efficiency
Color Scale
Min = 2.76e8
Max = 7.62e8
B
-DOX
+DOX
Mice ID
2 5 8 9 12 13 14 17 18 19 20 22 24
CAMKK2
p-AMPK(T172)
AMPK
GAPDH

### Figure S4

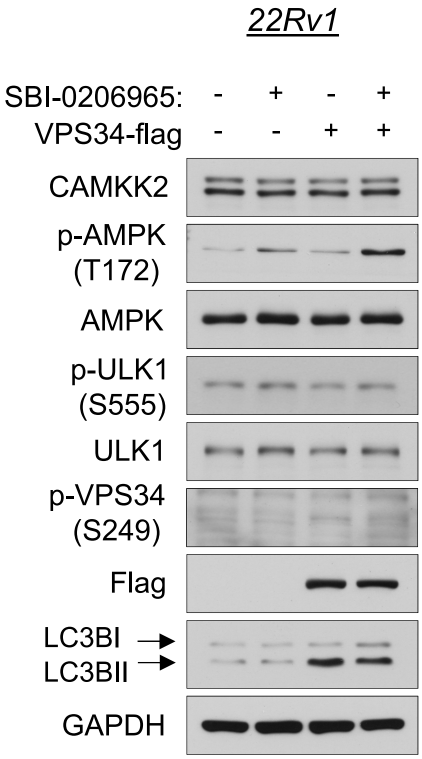
22Rv1
SBI-0206965: - + - +
VPS34-flag - - + +
CAMKK2
p-AMPK (T172)
AMPK
p-ULK1 (S555)
ULK1
p-VPS34 (S249)
Flag
LC3BI
LC3BII
GAPDH

Supplemental Table 1. siRNA, shRNA and sgRNA sequences.

| | Sequence |
|---|---|
| siRNA | |
| AMPK (*PRKAA1*) #1 | 5’-CACAGAAGGAUUUAAAUAUUGAGGG-3’ |
| AMPK (*PRKAA1*) #2 | 5’-ACCAUGAUUGAUGAUGAAGCCUUAA-3’ |
| AMPK (*PRKAA1*) #3 | 5’-UUAAGGCUUCAUCAUCAAUCAUGGU-3’ |
| shRNA | |
| *CAMKK2* | 5’-GGCATCGAGTACTTACACT-3’ |
| sgRNA | |
| *CAMKK2* | 5’-TGGAAGGTTTGATGTCACGG-3’ |
